# Structural analysis of *Helicobacter pylori* glutamate racemase in a monoclinic crystal form

**DOI:** 10.64898/2026.04.02.716094

**Authors:** Maria Spiliopoulou, Eike C. Schulz

## Abstract

Glutamate racemase (MurI) catalyzes the stereochemical interconversion of L-glutamate to D-glutamate, a key element of bacterial peptidoglycan biosynthesis. In this study, we present the crystal structure of *Helicobacter pylori* glutamate racemase at 1.43 Å and in monoclinic symmetry, as previously reported models, but different unit-cell parameters. The present model contains a single dimer and retains the previously described head-to-head dimer arrangement. The differences between the models arise from variations in unit-cell parameters, which lead to altered crystal packing interactions rather than changes in the quaternary assembly. The monomeric fold and active-site architecture remain conserved and are consistent with the catalytic features described for bacterial glutamate racemases. This structure provides an updated, high-resolution structural model for *H. pylori* glutamate racemase and highlights the variability in crystal packing within the same space group.

## 1 Introduction

Glutamate racemase (MurI) is a cofactor-independent enzyme that catalyzes the reversible stereochemical conversion of L-glutamate to D-glutamate, an essential component of bacterial peptidoglycan (murein). The production of D-glutamate is required for cell-wall biosynthesis, and inhibition of MurI disrupts peptidoglycan assembly, making the enzyme an attractive target for antibacterial drug development^1,2^. In addition to its metabolic role, MurI has been reported to exhibit secondary functions, such as inhibition of DNA gyrase, highlighting its functional versatility ^3^.

MurI enzymes catalyze racemization through a cofactor-independent two-base mechanism involving deprotonation of the substrate at the *α*-carbon followed by reprotonation on the opposite face. This reaction is mediated by conserved cysteine residues that act as general acid–base catalysts and MurI proteins typically consist of two domains (*α, β*) that form an active site at their interface, with catalytic residues positioned to facilitate stereoinversion of the bound substrate ^1,4^.

Glutamate racemase from *Helicobacter pylori* adopts the standard MurI folding and functions as a homodimer, which represents the biologically active form of the enzyme. The dimer interface contributes to enzyme stability and plays a role in shaping the active site environment as a head-to-head shape ^3,5^. Recent studies have further demonstrated that dimerization is functionally coupled to enzyme dynamics, with inter-subunit communication playing a critical role in catalysis. In particular, the identification of a cryptic allosteric pocket in *H. pylori* glutamate racemase has revealed that binding of small-molecule inhibitors can disrupt coupled motions between monomers, thereby impairing catalytic activity ^6^. These findings highlight the importance of dynamic interactions across the dimer interface and provide a mechanistic basis for allosteric regulation of MurI. Previous structural studies of *H. pylori* MurI have provided insight into substrate binding, catalytic mechanism and inhibition, consistently reporting dimeric assemblies in the crystal structures ^3,5^.

In this study, we present a crystallographic analysis of *H. pylori* glutamate racemase contributing to the further characterization of its molecular organization and dimer interface. Moreover, time-resolved crystallography (TRX) benefits from highly homogeneous crystals with sufficient solvent content to facilitate ligand diffusion. In this context, crystallization conditions for *Helicobacter pylori* glutamate racemase were explored to identify crystal forms with improved properties. Of particular interest is the protein dimer arrangement within the crystal lattice. This work provides additional structural information that contributes to understanding the variability in crystal packing and unit-cell organization in MurI enzymes.

## 2 Results and discussion

### 2.1 Crystal quality optimization

Crystallization conditions were initially based on previously reported conditions for *Helicobacter pylori* glutamate racemase, consisting of 0.1 M Tris pH 8.5, 0.2 M MgSO_4_, and 25% (w/v) PEG4000^5^. This condition was subsequently screened, with particular emphasis on varying pH (7.0 to 8.5) as well as MgSO_4_ (0.1-0.4 M) and PEG4000 (10-25%) concentrations. Adjustments of these parameters were performed to improve crystal quality, leading to the identification of conditions that yielded well-diffracting crystals.

Interestingly, crystal morphology was strongly affected by pH and PEG concentration variations (**Table 1**), regardless of the MgSO_4_ concentration. Low pH (<8.0) and low PEG (<20 %) conditions yielded long needles while the combination of higher pH values (≥ 8.0) and higher PEG (≥ 20%) concentration resulted in shorter, thicker needles, accompanied by altered unit-cell parameters.

**Table 1.**
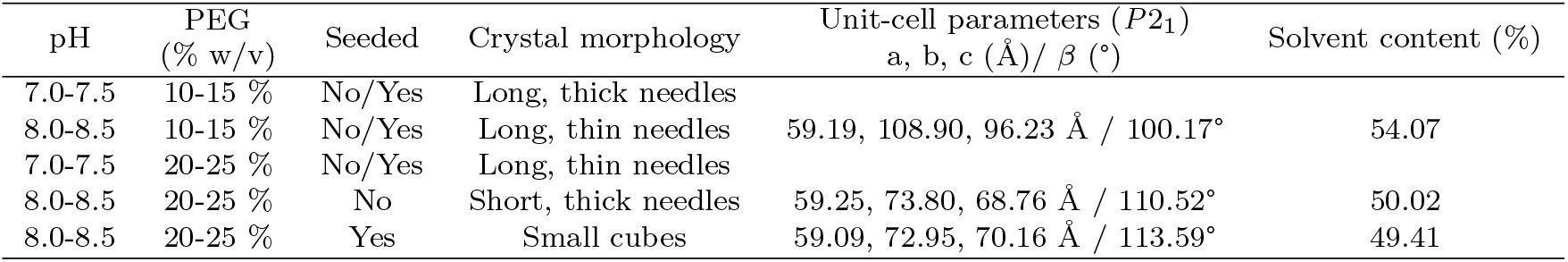
Summary of crystal morphologies and unit-cell observations for *H. pylori* MurI.

#### 2.1.1 Seeding

To further explore the influence of microseeding on crystal growth in sitting-drops, a seed stock was prepared from the short-thick needles, which were more stable and did not fragment during manipulation. This seed stock was then introduced into selected conditions from the full screen of varying salt/PEG concentration, and pH (mixing 1 *µ*L of 10 mg/mL protein, 0.8 *µ*L crystallization buffer and 0.2 *µ*L seeds). Under high pH and high PEG conditions, seeding produced crystals with a markedly different morphology (**Fig. 1**): the crystals were more homogeneous in both shape and size, while maintaining the same unit-cell parameters as the original short, thick needles. These observations indicate that seeding can stabilize crystal growth under specific conditions, leading to improved uniformity without altering the lattice parameters. This could be of particular interest, especially for expanding this study and using these crystals in TRX experiments^**?**^.

**Fig. 1.**
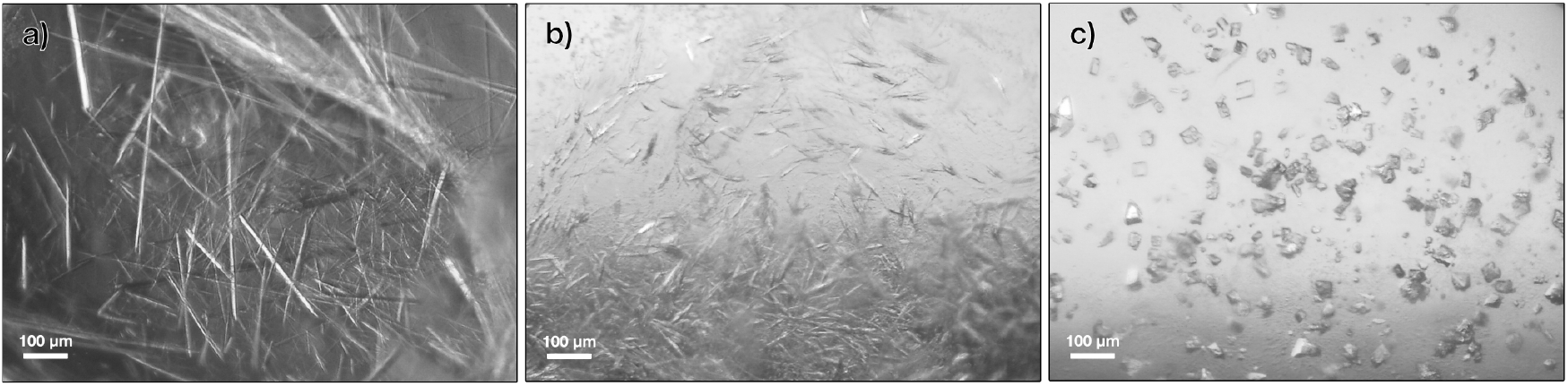
Photos of MurI protein crystals under different crystallization conditions. **(a)** 0.4 M MgSO_4_, 10% PEG, 0.1 M Tris pH 8.5; **(b)** 0.1 M MgSO_4_, 20% PEG, 0.1 M Tris pH 8.5; **(c)** 0.1 M MgSO_4_, 20% PEG,0.1 M Tris pH 8.5, seeding.

### 2.2 The monoclinic structural model of MurI

The crystal structure of *Helicobacter pylori* glutamate racemase was determined in a monoclinic space group, with unit-cell parameters of a = 59.09 Å, b = 72.95 Å, c = 70.16 Å and *β* = 113.59°. The asymmetric unit contains one homodimer, consistent with previously reported structures^5^. The overall fold of each monomer corresponds to the characteristic architecture of glutamate racemases, consisting of two domains (*α* and *β*) and the dimer adopts the conserved head-to-head arrangement (**Fig. 2**).

**Fig. 2.**
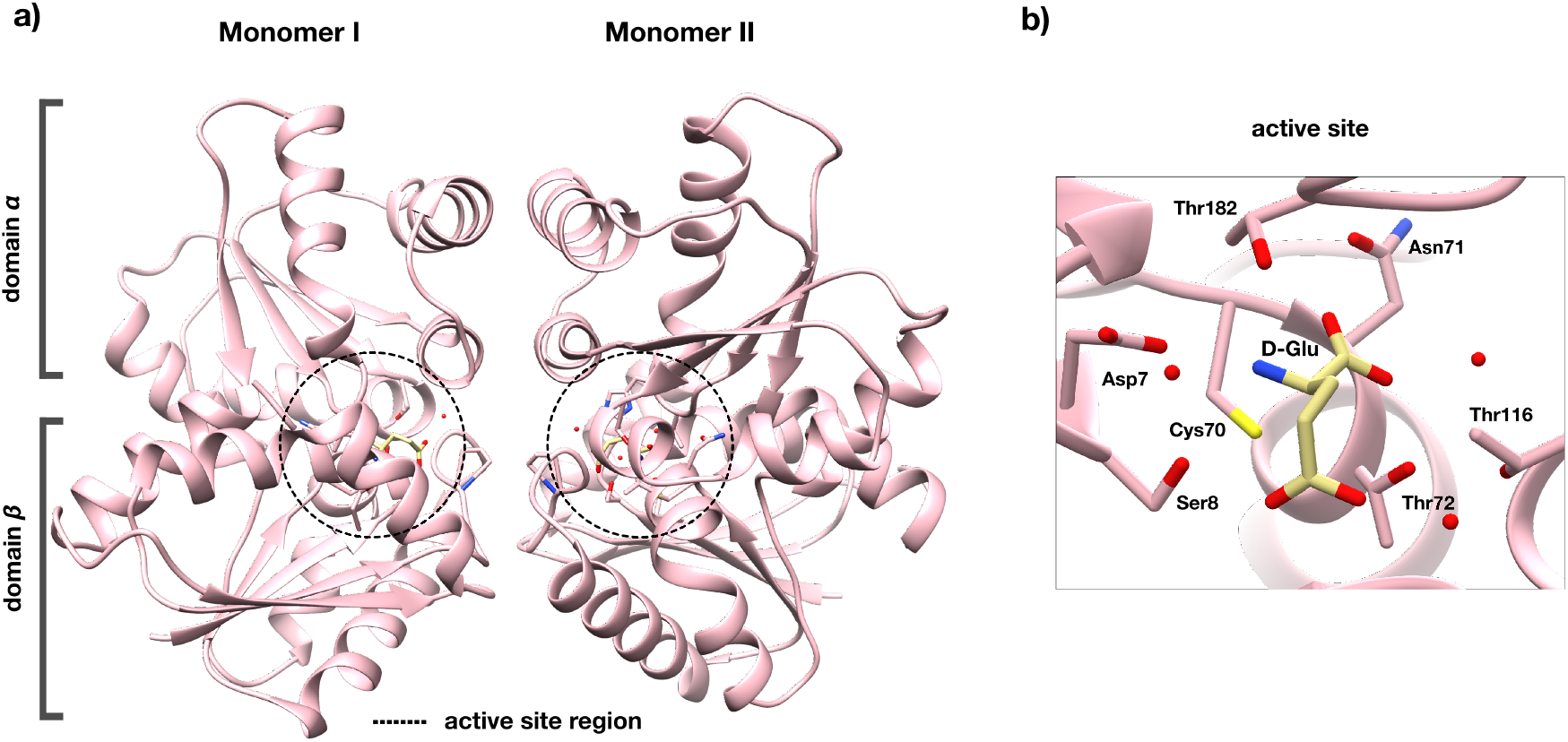
Cartoon representation of the MurI crystal structure. **(a)** The dimer arrangement and the active site regions indicated in circles. **(b)** Closeup of the active site and the surrounding amino acids.

The data extend to a resolution of 1.43 Å, resulting in a clear electron-density map and precise modeling of the overall protein and D-glutamate in both active sites. The ligand is well positioned within the catalytic pocket and interacts with the surrounding amino acids (**Fig. 3**). No major conformational differences are observed compared with earlier models, thus the refined structure represents a typical example of the enzyme in its dimeric, substrate-bound form.

**Fig. 3.**
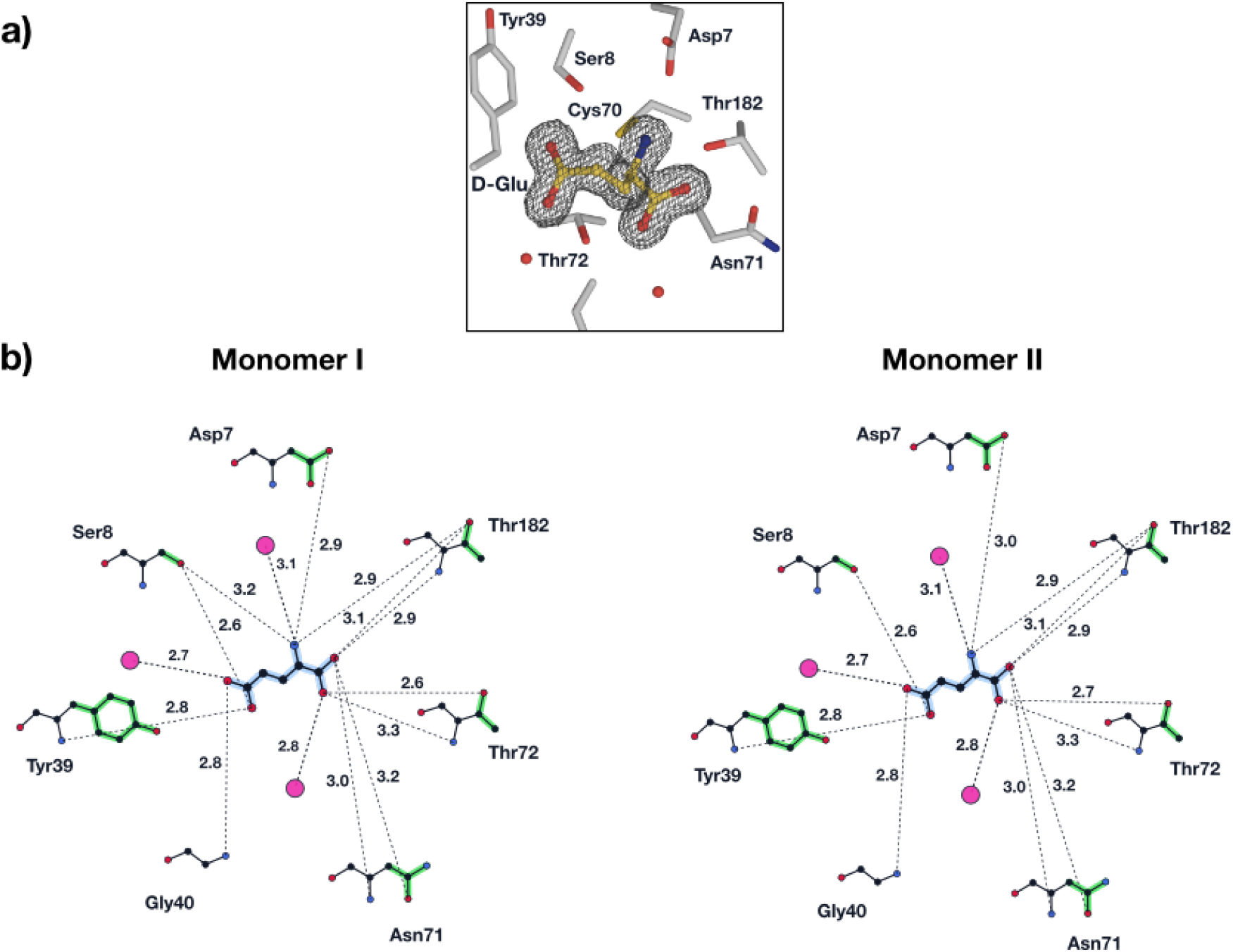
Molecular interactions of D-glutamate. **(a)** 3D representation of D-glutamate and the amino acids in the active site. Absolute END-RAPID map ^7^ contoured at 1.0 r.m.s.d. is represented by a grey mesh. **(b)** 2D representation of the hydrogen bonds that each D-glutamate molecule in the MurI dimer forms with the surrounding amino acids and water molecules. D-glutamate: center, highlighted in blue; H-bonds: dashed lines with lengths marked; amino acids: marked in 3-letter code, side-chains highlighted in green; water molecules: pink circles.

### 2.3 Crystal packing and symmetry-related interfaces

Structural comparison with previously reported models of *H. pylori* glutamate racemase reveals differences in dimer organization within the same crystallographic space group (*P* 2_1_). The deposited structures (PDB IDs: 2W4I and 2JFY) both adopt the head-to-head dimer arrangement, in which the active sites are oriented towards the internal face of the dimer interface ^5^.

Despite differences in crystallization conditions and asymmetric unit composition, both structures exhibit a similar mode of subunit association. The 2W4I model ^8^ contains two dimers per asymmetric unit, whereas 2JFY ^5^ contains a single dimer.

Our refined model (PDB ID: 29PA) was evaluated against the deposited structure 2JFY, as they appear to have similar asymmetric unit architecture, containing one dimer of MurI. Least-squares superposition yielded a C*α* root mean square deviation (RMSD) of 0.49 Å ^9,10^. Although our refined MurI structure crystallizes in the same space group, it appears to have different unit-cell parameters (a = 59.09 Å, b = 72.95, Å c = 70.16 Å, *β* = 113.59°, solvent content 49.41 %) compared with 2JFY (a = 52.28 Å, b = 78.96 Å, c = 59.14 Å, *β* = 92.64°, solvent content 42.50 %). The solvent-accessible surface area, measured using `PyMOL` ^11^, is 22815.32 Å^2^ for 29PA and 22387.01 Å^2^ for 2JFY.

Differences are observed in the crystal packing of symmetry-related molecules: 12 symmetry-related molecules are present in the 2JFY lattice, whereas only 8 are observed in the present structure. While the overall arrangement of molecules is largely preserved, variations in unit-cell affect the packing environment and the network of intermolecular interfaces within the crystal lattice (**Fig. 4**).

**Fig. 4.**
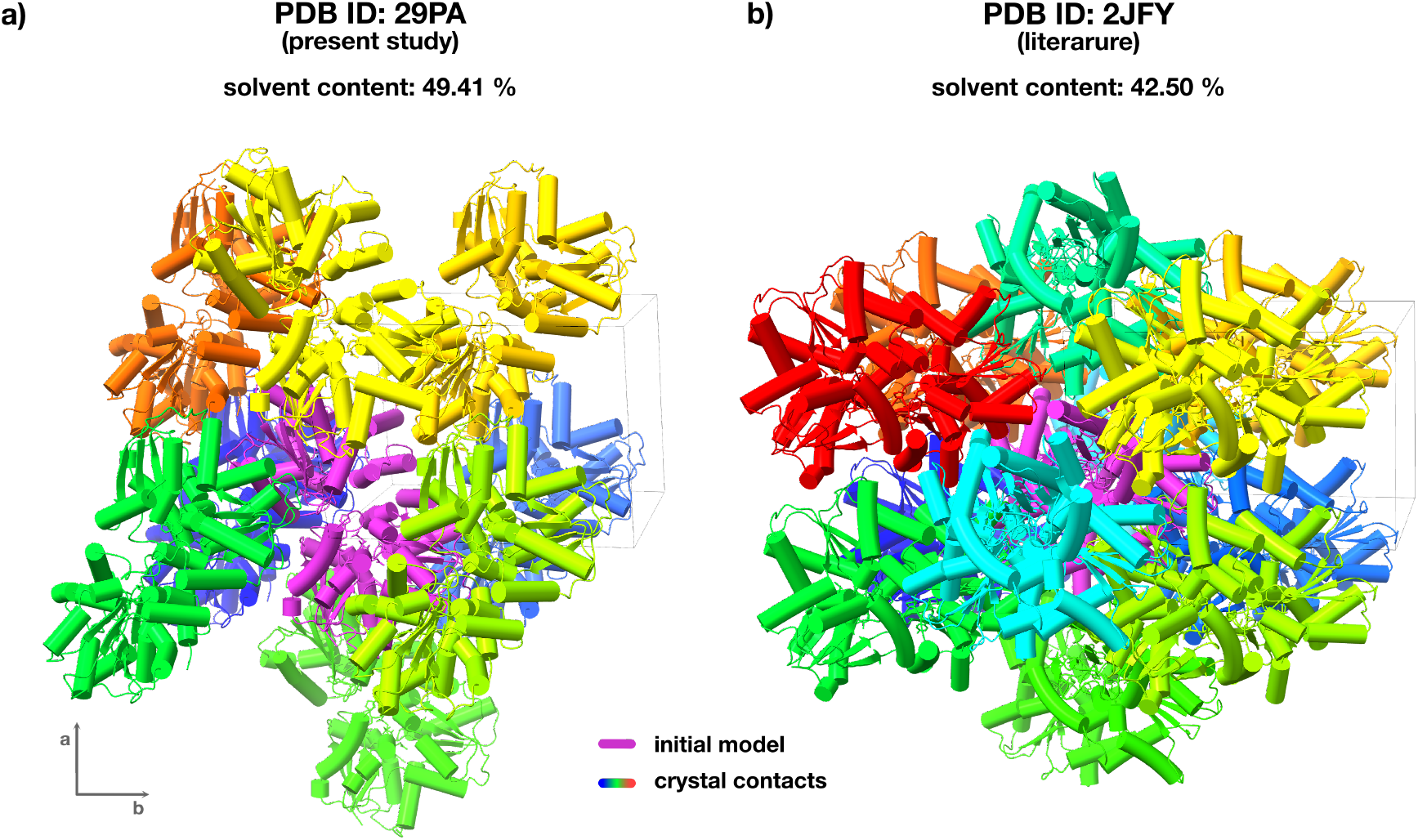
Crystal packing comparison. Symmetry related molecules that lead to crystal packing interactions for 29PA **(a)** and 2JFY **(b)**. The initial molecule in the assymetric unit is indicated in purple and the symmetry related ones in the rainbow pallette.

Crystal packing interactions were analysed using `PISA` ^12,13^ for the protein structure 2JFY in comparison with the refined model (this study, PDB ID: 29PA). The interface properties of the two structures were compared, focusing on symmetry-related contacts within the crystal lattice.

The comparative study revealed that the primary packing interaction (x,y,z) is highly conserved, with similar buried surface areas of approximately 1000 Å^2^ in both structures, indicating a shared dominant crystal contact. Consistent with this, several symmetry operators are common to both models, demonstrating that the overall lattice framework is preserved, despite differences in unit-cell parameters within the same space group.

In addition to this conserved primary interface, differences are observed in secondary packing interactions (**Table 2**). While the interface defined by the operator 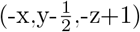 shows moderate similarity between the two structures, other contacts differ more substantially in buried surface area, particularly those generated by the operators (x-1,y,z) and 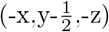. Furthermore, in PDB 2JFY most symmetry operators give rise to two equivalent interfaces, whereas in the present structure several operators are represented only once. These variations are reflected in differences in the extent and distribution of intermolecular contacts across the lattice. These variations are reflected in differences in the extent and distribution of intermolecular contacts across the lattice. Overall, the analysis indicates that, although our refined model (PDB ID: 29PA) and 2JFY share a conserved primary packing interaction and similar symmetry relationships, their crystal packing is not identical but instead represents a related arrangement with variations in secondary contacts.

**Table 2.**
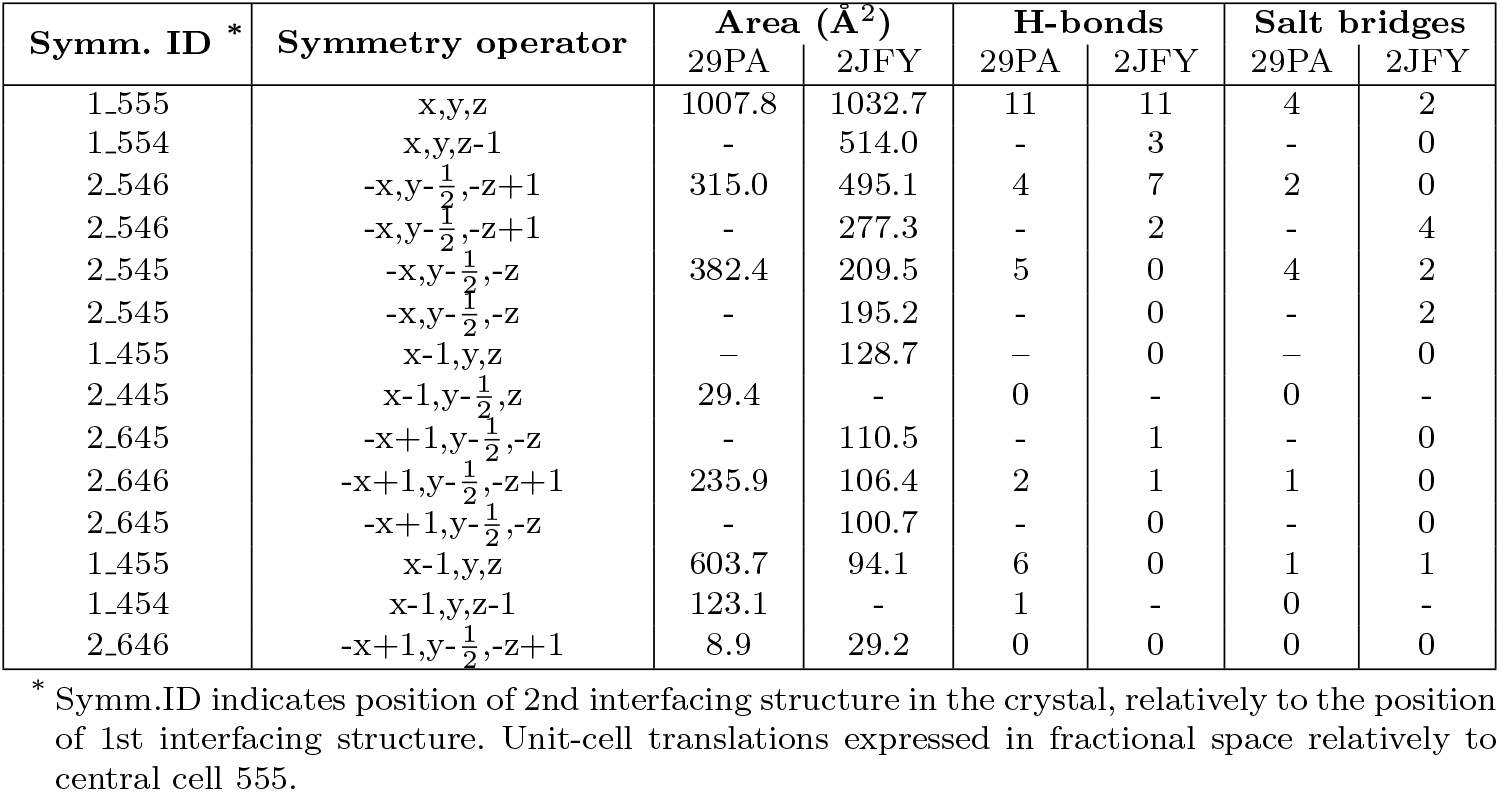
Comparison of crystal packing interfaces between the present structure and PDB 2JFY as calculated by PISA.

## 3 Conclusions

In the present study, the crystal structure of *H. pylori* MurI is presented in monoclinic symmetry, confirming the dimeric assembly in the asymmetric unit and the conserved active site architecture, in comparison with previously reported models for the same species. However, variations in the unit-cell parameters result in different crystal packing interactions; primary packing contacts within the crystal lattice are preserved, while secondary interfaces can differ within the same space group, as it is indicated by the comparative analysis.

The crystallization screening yielded crystal forms with distinct packing properties, reflecting differences in solvent content and lattice organization. Our observations highlight the influence of crystallization conditions on crystal packing and structural variability, with seeding contributing to improved homogeneity and change of crystal morphology, while preserving the overall protein architecture. Such features may be advantageous for TRX applications by enabling ligand diffusion within the crystal.

## 4 Materials and methods

### 4.1 Protein purification and crystallization

*Helicobacter pylori* MurI gene was cloned in pET28a(+) vector (Genscript) and expressed in Arctic Express *E. coli* cells grown in LB medium supplemented with 20 *µ*g/mL gentamycin and 50 *µ*g/mL kanamycin for the overnight starting culture at 37°C. For the large scale culture LB medium was supplemented with 50 *µ*g/mL kanamycin and cells were incubated at 30°C for 3 h (OD600≈0.7). Protein expression was induced by addition of IPTG to a final concentration of 0.001 M and the cells were further incubated at 13°C for 24 hours. The cells were harvested by centrifugation (7500*×*g, 20 min, 4°C) and pellets were stored at −20°C until purification. The pellets were resuspended in lysis buffer (0.05 M Tris pH 8.0, 0.15 NaCl, 5 % (v/v) glycerol) and sonicated for lysis.

The protein was purified by affinity (Ni) chromatography using 0.05M Tris-HCl, 0.15 M NaCl, 0.001 M DTT, 0.002 M ATP-Mg^2+^, pH 8.0 supplemented with 0.005 M and 0.5 M imidazole pH 8.0 for the binding and the elusion buffer correspondingly. Size exclusion chromatography followed with a final buffer of 0.2 M CH_3_COONH_4_ pH 7.4, 0.005 M DL-Glutamate, 0.001 M DTT^5^.

Crystallization was performed in sitting drops by mixing 1 *µ*L of 10 mg/mL protein, 0.8 *µ*L crystallization buffer and 0.2 *µ*L seeds, against 100 *µ*L of reservoir consisting of 0.1 M Tris pH 8.5, 0.1 M MgSO_4_ and 20 % (w/v) PEG4000.

### 4.2 X-ray data-collection

Single crystal rotation datasets were collected at beamline P13 at PETRA-III (DESY, Hamburg) ^14^ at the EMBL unit Hamburg. Data were collected on an EIGER 16 M detector at an energy of 11.7 keV. The beam was focused to 70 × 70 µm with a flux 4.97×10^11^ ph/s. Data collection and structure refinement parameters are summarized in **Table 3**.

**Table 3.** Data collection and refinement statistics of the MurI data. *Values in the highest resolution shell are shown in parentheses*.

| Sample | MurI |
| --- | --- |
| PDB ID | 29PA |
| **Data collection** |  |
| Source | P13, EMBL |
| Beam size ($\mu\text{m}$) | 70 x 70 |
| Flux (ph/s) | $4.97 \times 10^{11}$ |
| Energy (keV) | 11.7 |
| Exposure time (ms) | 8 |
| Number of images | 3600 |
| Space group | $P2_1$ |
| Unit-cell parameters |  |
| a, b, c ($\text{\AA}$) | 59.09, 72.95, 70.16 |
| $\alpha, \beta, \gamma$ ($^\circ$) | 90, 113.59, 90 |
| Resolution range ($\text{\AA}$) | 64.29-1.43 (1.59-1.43) |
| Total reflections | 460372 (20886) |
| Unique reflections | 64959 (3248) |
| Mean $I/\sigma$ (I) | 9.1 (1.4) |
| Completeness (ellipsoidal) | 87.4 (40.9) |
| Multiplicity | 7.1 (6.4) |
| $CC_{1/2}$ | 0.998 (0.558) |
| **Refinement** |  |
| Resolution range ($\text{\AA}$) | 48.23-1.43 |
| Number of reflections | 64951 |
| $R_{\text{work}}$ | 15.49 |
| $R_{\text{free}}$ | 19.66 |
| Wilson $B$-factor ($\text{\AA}^2$) | 18.31 |
| Mean $B$-factor ($\text{\AA}^2$) | |
| Overall | 25.35 |
| Protein | 24.63 |
| Water | 33.96 |
| Ligands | 14.09 |
| **Model quality** |  |
| *RMS deviations* |  |
| Bond length ($\text{\AA}$) | 0.015 |
| Bond angles ($^\circ$) | 2.07 |

### 4.3 Data processing

Diffraction data were automatically processed with `autoproc` using `StarAniso`^15–18^. Molecular replacement was conducted in `phaser` using PDB-ID 2JFY as starting model. Structures were refined using iterative cycles of `refmac` ^19^ and `coot`^20^. Molecular images were generated in `PyMol` ^11^ and `UCSF Chimera` ^21^.

Molecular interactions were inferred from parsed PDB structures using distance heuristics and projected into a 2D plane, with residues spatially organized around the central ligand. The resulting visualizations were rendered as scalable vector graphics (SVG), depicting atoms, bonds, and labels, with dashed lines used to represent non-covalent interactions.

## Acknowledgments

We would like to thank G.Gore for help with the 2D-ligand interaction diagrams. ES acknowledges support by the Federal Ministry of Education and Research, Germany, under grant number 01KI2114. Funded by the European Union (ERC, DynaPLIX, SyG-2022 101071843). Views and opinions expressed are however those of the author(s) only and do not necessarily reflect those of the European Union or the European Research Council Executive Agency (ERCEA). Neither the European Union nor the granting authority can be held responsible for them.

## Data and materials availability

The data that support this study are available from the corresponding authors upon request. All crystallographic data have been deposited in the Protein Data Bank (PDB) under following accession number: 29PA. Further details are available in **Table 3**.

## References

[1] Grishin, A. V. et al. Resistance to peptidoglycan-degrading enzymes. Critical Reviews in Microbiology 46, 703–726 (2020).

[2] Keating, T. A. Resistance mechanism to an uncompetitive inhibitor of a single-substrate, single-product enzyme: A study of helicobacter pylori glutamate racemase. Future Medicinal Chemistry 5, 1203–1214 (2013).

[3] Fisher, S. L. Glutamate racemase as a target for drug discovery. Microbial Biotechnology 1, 345–360 (2008).

[4] Ruzheinikov, S. N., Taal, M. A., Sedelnikova, S. E., Baker, P. J. & Rice, D. W. Substrate-induced conformational changes in bacillus subtilis glutamate racemase and their implications for drug discovery. Structure 13, 1707–1713 (2005).

[5] Lundqvist, T. et al. Exploitation of structural and regulatory diversity in glutamate racemases. Nature 447, 817–822 (2007).

[6] Chheda, P. R. et al. Decrypting a cryptic allosteric pocket in h. pylori glutamate racemase. Communications Chemistry 4 (2021).

[7] Lang, P. T., Holton, J. M., Fraser, J. S. & Alber, T. Protein structural ensembles are revealed by redefining X-ray electron density noise. Proc. Natl. Acad. Sci. U. S. A. 111, 237–242 (2014).

[8] Geng, B. et al. Potent and selective inhibitors of helicobacter pylori glutamate racemase (muri): Pyridodiazepine amines. Bioorganic amp; Medicinal Chemistry Letters 19, 930–936 (2009).

[9] Kabsch, W. A solution for the best rotation to relate two sets of vectors. Acta Crystallographica Section A 32, 922–923 (1976).

[10] Carugo, O. & Pongor, S. A normalized root-mean-spuare distance for comparing protein three-dimensional structures. Protein Science 10, 1470–1473 (2001).

[11] Schrodinger, L. The pymol molecular graphics system. Version 1, 8 (2015).

[12] Krissinel, E. & Henrick, K. Inference of macromolecular assemblies from crystalline state. Journal of Molecular Biology 372, 774–797 (2007).

[13] Baskaran, K., Duarte, J. M., Biyani, N., Bliven, S. & Capitani, G. A pdb-wide, evolution-based assessment of protein-protein interfaces. BMC Structural Biology 14 (2014).

[14] Cianci, M. et al. P13, the embl macromolecular crystallography beamline at the low-emittance petra iii ring for high- and low-energy phasing with variable beam focusing. Journal of Synchrotron Radiation 24, 323–332 (2017).

[15] Vonrhein, C. et al. Data processing and analysis with the autoproc toolbox. Acta Crystallographica Section D: Biological Crystallography 67, 293–302 (2011).

[16] Tickle, I. et al. Staraniso global phasing ltd. Cambridge, United Kingdom (2018).

[17] Kabsch, W. xds. Acta Crystallographica Section D: Biological Crystallography 66, 125–132 (2010).

[18] Agirre, J. et al. The ccp4 suite: integrative software for macromolecular crystallography. Acta Crystallographica Section D: Structural Biology 79, 449–461 (2023).

[19] Vagin, A. A. et al. Refmac5 dictionary: organization of prior chemical knowledge and guidelines for its use. Acta Crystallographica Section D Biological Crystallography 60, 2184–2195 (2004).

[20] Emsley, P., Lohkamp, B., Scott, W. G. & Cowtan, K. Features and development of coot. Acta Crystallographica Section D: Biological Crystallography 66, 486–501 (2010).

[21] Pettersen, E. F. et al. Ucsf chimera—a visualization system for exploratory research and analysis. Journal of Computational Chemistry 25, 1605–1612 (2004).

